# Dynamic regulation of RNA Polymerase III transcription in mouse embryonic stem cells during heat shock stress

**DOI:** 10.1101/2024.11.28.625959

**Authors:** Thomas F. Nguyen, James Z.J. Kwan, Jennifer E. Mitchell, Jieying H. Cui, Sheila S. Teves

## Abstract

Cells respond to many different types of stresses by overhauling gene expression patterns, both at the transcriptional and translational level. Under heat stress, global transcription and translation are inhibited, while the expression of chaperone proteins are preferentially favored. As the direct link between mRNA transcription and protein translation, tRNA expression is intricately regulated during the stress response. Despite extensive research into the heat shock response (HSR), the regulation of tRNA expression by RNA Polymerase III (Pol III) transcription has yet to be fully elucidated in mammalian cells. Here, we examine the regulation of Pol III transcription during different stages of heat shock stress in mouse embryonic stem cells (mESCs). We observe that Pol III transcription is downregulated after 30 minutes of heat shock, followed by an overall increase in transcription after 60 minutes of heat shock. This effect is more evident in tRNAs, though other Pol III gene targets are also similarly affected. Notably, we show that the downregulation at 30 minutes of heat shock is independent of HSF1, the master transcription factor of the HSR, but that the subsequent increase in expression at 60 minutes requires HSF1. Taken together, these results demonstrate an adaptive RNA Pol III response to heat stress, and an intricate relationship between the canonical HSR and tRNA expression.

**Article Summary:** This study explores the regulation of RNA Polymerase III (Pol III) transcription during heat shock in mouse embryonic stem cells (mESCs). Results show that tRNA transcription is downregulated after 30 minutes of heat shock, but increases after 60 minutes, while other Pol III targets remain unaffected. Importantly, the initial downregulation is independent of heat shock factor 1 (HSF1), the key regulator of the heat shock response, but the subsequent increase in tRNA expression depends on HSF1. These findings reveal an adaptive mechanism of Pol III activity under heat stress, highlighting a complex interplay between heat shock response and tRNA expression.

## Introduction

Cells are often exposed to various environmental stressors that necessitate rapid and coordinated responses at the molecular level, including the reprogramming of gene expression (Fulda et al. 2010). Among the best-characterized stress responses is the heat shock response (HSR), governed by the master regulator Heat Shock Factor 1 (HSF1) (Ritossa 1962; Lindquist 1986; Fulda et al. 2010). Upon heat shock, HSF1 is activated and induces the rapid transcription of heat shock protein (HSPs) genes by RNA Polymerase II (Pol II) (Tissiéres et al. 1974; Lindquist and Craig 1988; Åkerfelt et al. 2010; Björk and Sistonen 2010). These HSPs function as molecular chaperones, refolding misfolded proteins during heat stress to maintain proteostasis (Lindquist and Craig 1988; Richter et al. 2010). Simultaneously, global transcription and translation are largely inhibited to allocate cellular resources properly during heat stress (Mahat et al. 2016; Aprile-Garcia et al. 2019; Pessa et al. 2024).

Although most of the research on stress responses has focused on the transcriptional regulation of mRNA and stress-induced genes, the regulation of transfer RNAs (tRNAs) during stress remains less understood. As critical links between transcription and translation, the proper regulation of tRNA expression under stress conditions is essential for maintaining proteostasis. RNA Polymerase III (Pol III) transcribes specific noncoding genes, including tRNAs, 5S rRNA and U6 snRNA (Weinmann and Roeder 1974; Ullu and Weiner 1985; Krüger and Benecke 1987; Reddy et al. 1987). Studies in yeast and human cells show that stress conditions, including oxidative stress and serum starvation, lead to a global reduction in tRNA abundance (Desai et al. 2005; Michels et al. 2010; Orioli et al. 2016; Torrent et al. 2018). Stress can also disrupt tRNA maturation and function, as observed with arsenite and methyl methanesulfonate in yeast (Yoluç et al. 2021). Intriguingly, tRNA nuclear relocalization has been proposed as a mechanism to downregulate translation during stress in both yeast and mammalian cells (Shaheen and Hopper 2005; Miyagawa et al. 2012). However, how Pol III regulation adapts to heat stress over time in mammalian cells, and whether the canonical HSR is involved, remains to be fully elucidated.

In this study, we focused on the regulation of Pol III-driven tRNA transcription during heat shock stress in mouse embryonic stem cells (mESCs). We aimed to determine how Pol III activity and tRNA transcription are modulated at different stages of heat shock and whether these changes are coordinated with the canonical heat shock response as governed by heat shock factor 1 (HSF1). Our findings reveal an intricate regulation of Pol III during heat stress, with tRNA transcription showing distinct temporal dynamics, and highlight the role of HSF1 in modulating these processes.

## Results

### Transcription of tRNA genes is dynamically regulated during heat shock

We explored how tRNA transcription adapts to heat stress over time by exposing mESCs to heat shock (HS) at 42°C for 30 and 60 minutes (HS30, HS60), and measuring Pol III occupancy using Cleavage Under Targets and Tagmentation (CUT&Tag), a high-resolution chromatin profiling technique (Kaya-Okur et al. 2019), in two biological replicates for each condition (Figure S1A). Reads were aligned to the mouse genome, normalized using ChIP-seq-Spike-In-Free method (Jin et al. 2020), and replicates were merged. Gene browser tracks at specific tRNA gene loci show Pol III binding decreased 2-fold upon 30 minutes of HS (Figure 1A) compared to unstressed (HS0) conditions, consistent with previous findings using different stressors in other species (Desai et al. 2005; Michels et al. 2010; Orioli et al. 2016). Surprisingly, Pol III binding returned back to normal under continued HS for 60 minutes (Figure 1A). A similar trend was observed for all tRNA genes when we plotted the Pol III occupancy in a 400 bp window surrounding the transcription start site (TSS) for all tRNA genes (Figure 1B).

**Figure 1.**
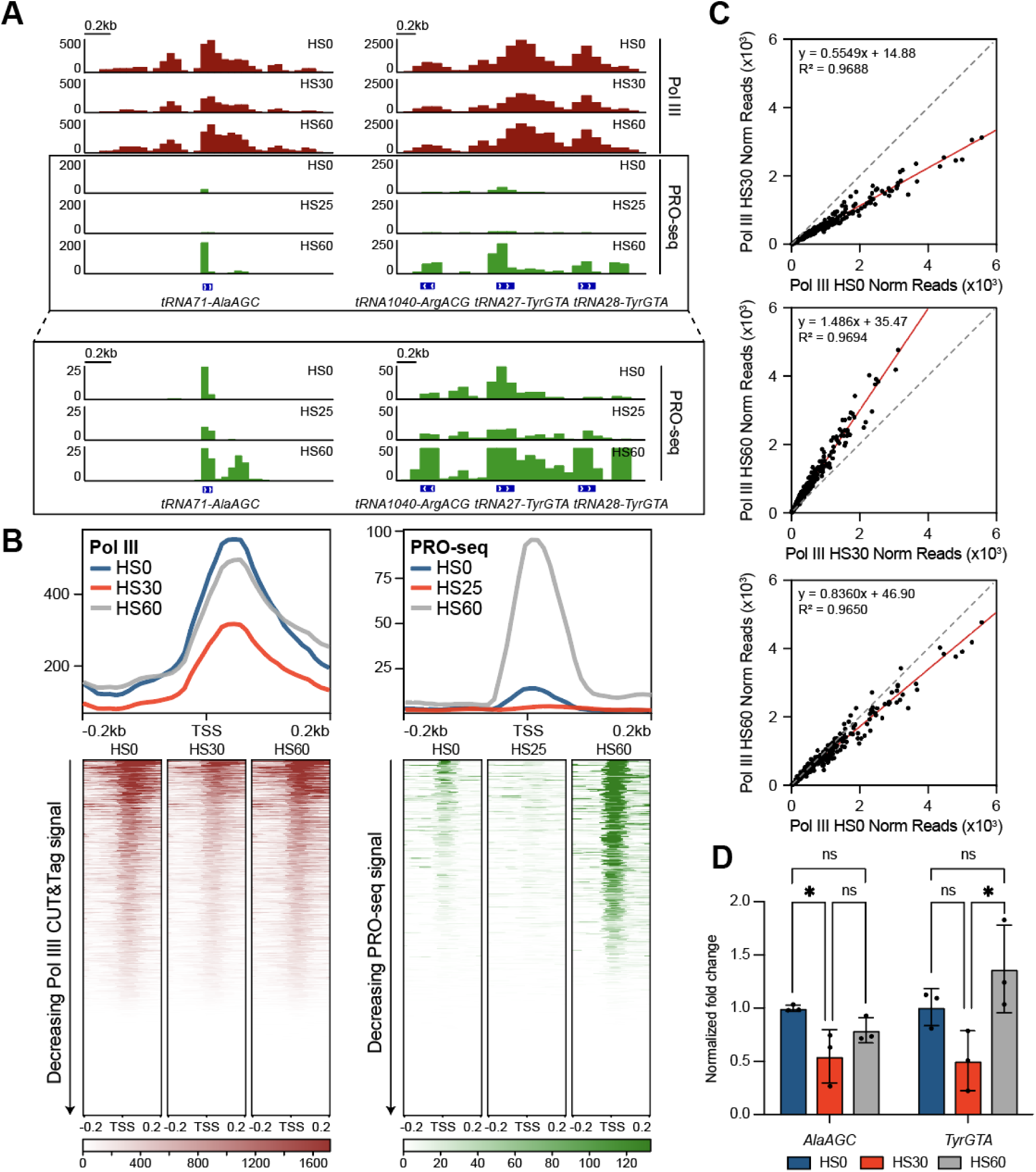
Transcription of tRNAs is dynamically regulated during different stages of heat shock. **(A)** Gene browser tracks at *tRNA71-AlaAGC* (left) and *tRNA1040-ArgACG, tRNA27-TyrGTA, tRNA28-TyrGTA* (right) of Pol III CUT&Tag (maroon, n=2) in unstressed mESCs (HS0) or with 30 minutes (HS30) and 60 minutes (HS60) of heat shock treatment, and PRO-seq (green, from Vihervaara et al., 2021, n=2) in unstressed wildtype MEFs or with 25 and 60 minutes of heat shock (HS25, HS60 respectively). PRO-seq tracks with adjusted y-axis are shown below within the boxed regions. (**B)** Genome-wide average plots (top) and heatmaps (bottom) arranged by decreasing signal for Pol III CUT&Tag in mESCs and PRO-seq in MEFs in a 400 bp window surrounding the TSS of all tRNA genes. **(C)** Normalized read counts of Pol III CUT&Tag signal in unstressed vs. HS30-treated mESCs (top), unstressed vs. HS60-treated mESCs (middle), and HS30-treated vs. HS60-treated mESCs (bottom) from the TSS to the TES of all tRNA genes. **(D)** BRI-qPCR analysis of *tRNA71-AlaAGC* and *tRNA27-TyrGTA* in HS0 (blue), HS30 (red), and HS60 (gray) treated mESCs normalized to NLuc signal (n=3, mean ± SD). Statistical analysis was performed using one-way ANOVA. ns: non-significant. *: p ≤ 0.05.

To quantify the change in Pol III binding over time under HS, we plotted the Pol III CUT&Tag normalized read count values on tRNAs as scatter plots, comparing HS0 versus HS30, HS30 versus HS60, and HS0 versus HS60 samples (Figure 1C). Linear regression analyses showed a ∼50% decrease in HS0 versus HS30, followed by an increase of ∼50% in HS30 versus HS60. The HS0 versus HS60 scatter plot confirmed the genome-wide trends. These analyses also show that both downregulation and subsequent recovery occur for all transcribed tRNAs, and not a unique subset. Importantly, immunoblotting of whole cell lysates for the RPC7 subunit of Pol III showed no changes in unstressed, HS30, and HS60 conditions, suggesting that the changes in Pol III occupancy during HS is not due to changes in Pol III protein levels (Figure S2A).

The CUT&Tag data showed changes in Pol III occupancy upon HS, but to directly assess Pol III activity, we performed several analyses. First, we re-analyzed previously published Native Elongating Transcripts coupled with sequencing (NET-seq) datasets in HS30-treated mESCs compared to unstressed controls (Kwan et al. 2023). NET-seq captures newly transcribed RNA from elongating RNA Pols at single nucleotide resolution through extensive fractionation (Mayer et al. 2015). We observed a 2-fold decrease in signal at *AlaAGC* and *TyrGTA* gene loci (Figure S2B) and genome-wide across all tRNAs (Figure S2C) in HS30-treated mESCs compared to unstressed mESCs. A scatterplot comparing normalized NET-seq reads from the TSS to the transcription end site (TES) of all tRNAs in unstressed versus HS30-treated mESCs also showed a similar decrease of signal in HS30-treated cells (Figure S2D), corroborating the Pol III CUT&Tag data that tRNA transcription is decreased in mESCs after 30 minutes of HS.

Secondly, we re-analyzed previously published Precision Run-On-sequencing (PRO-seq) datasets (Vihervaara et al. 2021) of mouse embryonic fibroblasts (MEFs) treated with 25 minutes and 60 minutes of HS (HS25 and HS60, respectively). Similar to NET-seq, PRO-seq maps the active site of RNA Pols with single base resolution, providing a direct measure of transcriptional activity (Mahat et al. 2016). Consistent with the CUT&Tag and NET-seq analyses, we observed a 2-3 fold decrease in PRO-seq signal at *AlaAGC* and *TyrGTA* gene loci (Figure 1A) and genome-wide across all tRNAs (Figure 1B, Figure S1A) in HS25-treated MEFs compared to unstressed MEFs. We also observed a 5-8-fold increase in *AlaAGC* and *TyrGTA* levels (Figure 1A) and across all tRNAs genome-wide (Figure 1B, Figure S1A) in HS60-treated MEFs compared to HS25 treatment and unstressed cells. These analyses show that the dynamic HS-related changes in Pol III activity that we observed in mESCs also occur in MEFs.

Thirdly, we performed metabolic labeling with 5’-bromo-uridine (5-BrU) to directly measure changes in tRNA expression over 30 and 60 minutes of HS in mESCs (Figure S2E). 5-BrU-labeled nascent RNA was immunoprecipitated (Imamachi et al. 2014) and quantified by RT-qPCR (BRI-qPCR) in unstressed, HS30-treated, and HS60-treated mESCs (Figure 1D). In vitro transcribed 5-BrU labeled NanoLuc® Luciferase (NLuc) RNA was spiked-in for normalization (England et al. 2016). Analyzing two different tRNA genes (*AlaAGC, TyrGTA*), we observe a decreasing trend in nascent RNA after 30 minutes of HS, followed by an increasing trend after 60 minutes (Figure 1D). Altogether, these data indicate an adaptive and dynamic regulation of RNA Pol III transcription of tRNAs during HS.

### HSF1 regulates the adaptive recovery of tRNA transcription during heat shock

As the master regulator of the HSR, HSF1 induces the Pol II-mediated transcription of HSP genes (Åkerfelt et al. 2010; Björk and Sistonen 2010), but how this canonical HSR feedbacks into tRNA transcription is unknown. Previously generated *Hsf1*^-/-^ mESCs (Price et al. 2023) were exposed to HS at 42°C for 30 and 60 minutes. We confirmed HSF1 knockout by western blot analysis (Figure 2A), and that these cells failed to induce HSP genes, as measured by RT-qPCR of the *Hspa1a* gene (Figure S3A). To investigate how the chromatin occupancy of Pol III is affected upon HS treatment in the absence of HSF1, we performed Pol III CUT&Tag analysis in two biological replicates (Figure S3B) of unstressed, HS30-treated, and HS60-treated *Hsf1*^-/-^ mESCs, normalized using the ChIP-seq-Spike-In-Free method (Jin et al. 2020). Similar to wild type cells, gene browser tracks at specific tRNA loci show that Pol III is bound abundantly in unstressed conditions and decreases 2-fold after 30 minutes of HS (Figure 2B). In contrast, the observed recovery of Pol III occupancy after 60 minutes of HS in wild type cells was blunted in *Hsf1*^-/-^ mESCs (Figure 2B). We plotted the Pol III occupancy in a 400 bp window surrounding the TSS for all tRNA genes in unstressed, HS30, and HS60-treated *Hsf1*^-/-^ mESCs and observed a similar trend across all tRNA genes (Figure 2C).

**Figure 2.**
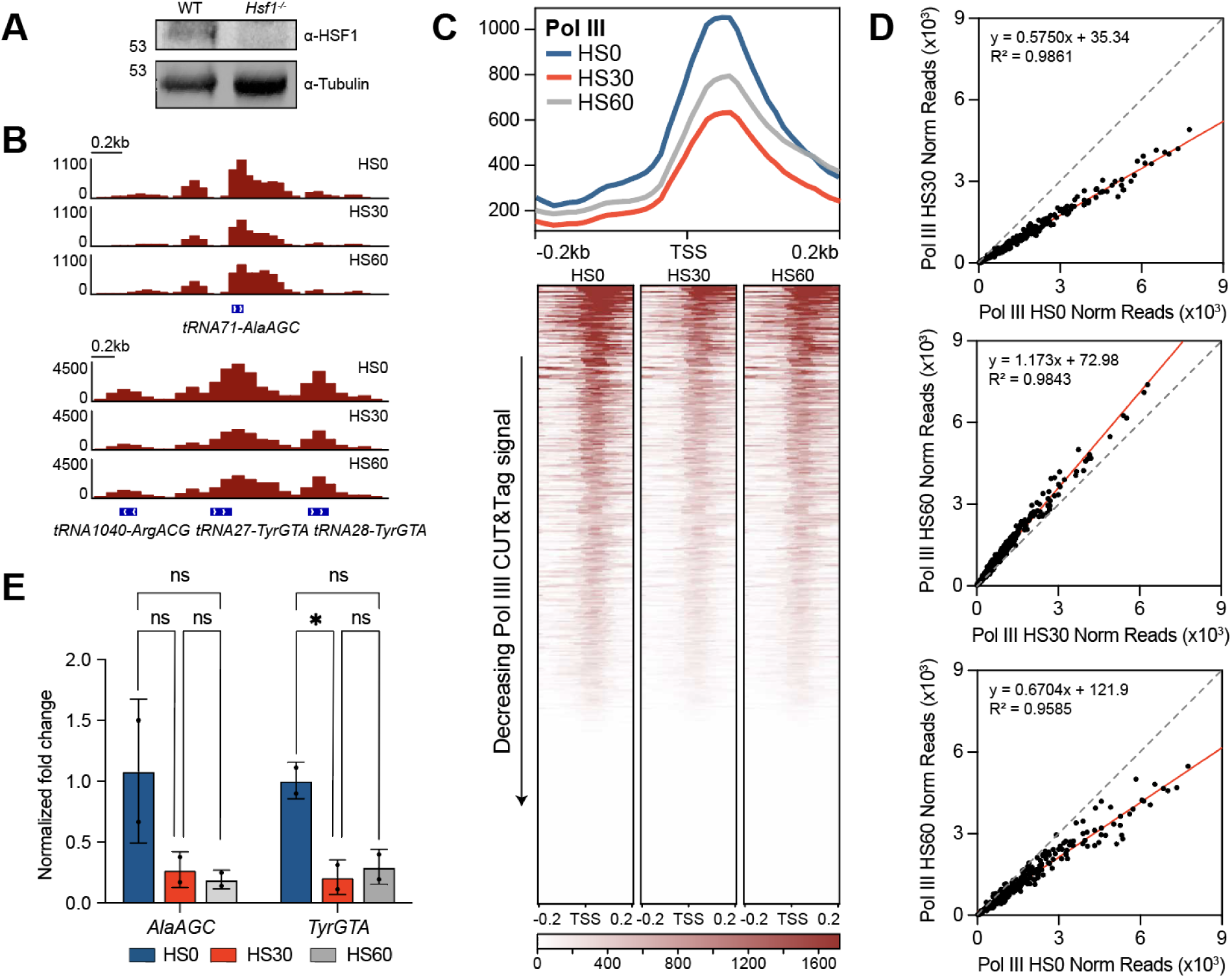
tRNA transcription regulation is partially dependent on HSF1. **(A)** Immunoblot analysis with α-HSF1 and α-Tubulin of wildtype (WT) and *Hsf1*^-/-^ mESC whole cell extracts. **(B)** Gene browser tracks at *tRNA71-AlaAGC* (left) and *tRNA1040-ArgACG, tRNA27-TyrGTA, tRNA28-TyrGTA* (right) of Pol III CUT&Tag (maroon, n=2) in unstressed *Hsf1*^-/-^ mESCs (HS0) or with 30 minutes (HS30) and 60 minutes (HS60) of heat shock treatment. **(C)** Genome-wide average plots (top) and heatmaps (bottom) for Pol III CUT&Tag signal in a 400 bp window surrounding the TSS of all tRNA gene, arranged by decreasing signal in *Hsf1*^-/-^ mESCs after HS0, HS30, and HS60 treatments. **(D)** Normalized read counts of Pol III CUT&Tag signal in unstressed vs. HS30-treated *Hsf1*^-/-^ mESCs (top), unstressed vs. HS60-treated *Hsf1*^-/-^ mESCs (middle), and HS30-treated vs. HS60-treated *Hsf1*^-/-^ mESCs (bottom) from the TSS to the TES of all tRNA genes. **(E)** BRI-qPCR analysis of *tRNA71-AlaAGC* and *tRNA27-TyrGTA* in HS0 (blue), HS30 (red), and HS60 (gray) treated *Hsf1*^-/-^ mESCs normalized to NLuc signal (n=2, mean ± SD). Statistical analysis was performed using one-way ANOVA. ns: non-significant. *: p ≤ 0.05.

To further quantify the changes in Pol III occupancy, we generated scatterplots with normalized read count values of Pol III on tRNAs in HS0 versus HS30-treated cells, HS30-treated versus HS60-treated, and in HS0 versus HS60-treated cells and performed linear regression analysis. We observed that most points fell below the diagonal, decreasing ∼45% in the HS0 versus HS30-treated samples, and increasing ∼15% in HS30-treated versus HS60-treated samples (Figure 2D). The HS0 versus HS60 scatterplot shows a ∼30% overall decrease, reflecting the stunted recovery observed between these conditions. As in wild type cells, western blot analysis of unstressed, HS30-treated, and HS60-treated *Hsf1*^-/-^ mESC whole cell lysates showed comparable RPC7 levels across all conditions, suggesting that HS treatment did not affect Pol III protein levels in *Hsf1*^-/-^ mESCs (Figure S3C).

We then tested whether Pol III occupancy is consistent with activity by performing BRI-qPCR in unstressed, HS30-treated, and HS60-treated *Hsf1*^-/-^ mESCs (Figure 2E). BRI-qPCR of the tRNA target *AlaAGC* showed an average decrease in signal in HS30-treated *Hsf1*^-/-^ mESCs compared to unstressed conditions and that this signal remained low in HS60-treated conditions (Figure 2E). However, BRI-qPCR of the tRNA target *TyrGTA* showed a significant decrease in RNA levels in HS30-treated mESCs compared to unstressed conditions and that this decrease in RNA levels persists into HS60-treated conditions (Figure 2E). Taken together, these results suggest that the downregulation of tRNAs during mid-HS is independent of HSF1, while the recovery of tRNA transcription during late HS depends on the presence of HSF1 in mESCs.

### HSF1 modulates the adaptive response of other Pol III gene classes

In addition to tRNA genes, Pol III also transcribes the 5S ribosomal RNA, a core component of the ribosome, 7S RNAs, and other small non-coding RNAs (Weinmann and Roeder 1974; Ullu and Weiner 1985; Krüger and Benecke 1987; Reddy et al. 1987). To investigate how these other classes of Pol III genes adapt to heat stress over time, we analyzed our Pol III CUT&Tag data at 42°C for HS30 and HS60 at the 5S, 7S1 and 7SK genes. Gene browser tracks at these gene loci show strong Pol III occupancy under unstressed conditions and a modest decrease upon 30 minutes of HS, although not to the degree as seen for tRNAs (Figure 3A). As with tRNAs, Pol III occupancy at the 5S, 7S1 and 7SK genes returned back to normal unstressed levels after 60 minutes of HS (Figure 3A, Figure S4A). We quantified these results by averaging the normalized read counts within gene bodies for each replicate, and the bar plots for these samples confirm this trend (Figure 3B). These results suggest that other classes of Pol III genes are dynamically regulated under HS stress, but the degree of regulation varies between these genes.

**Figure 3.**
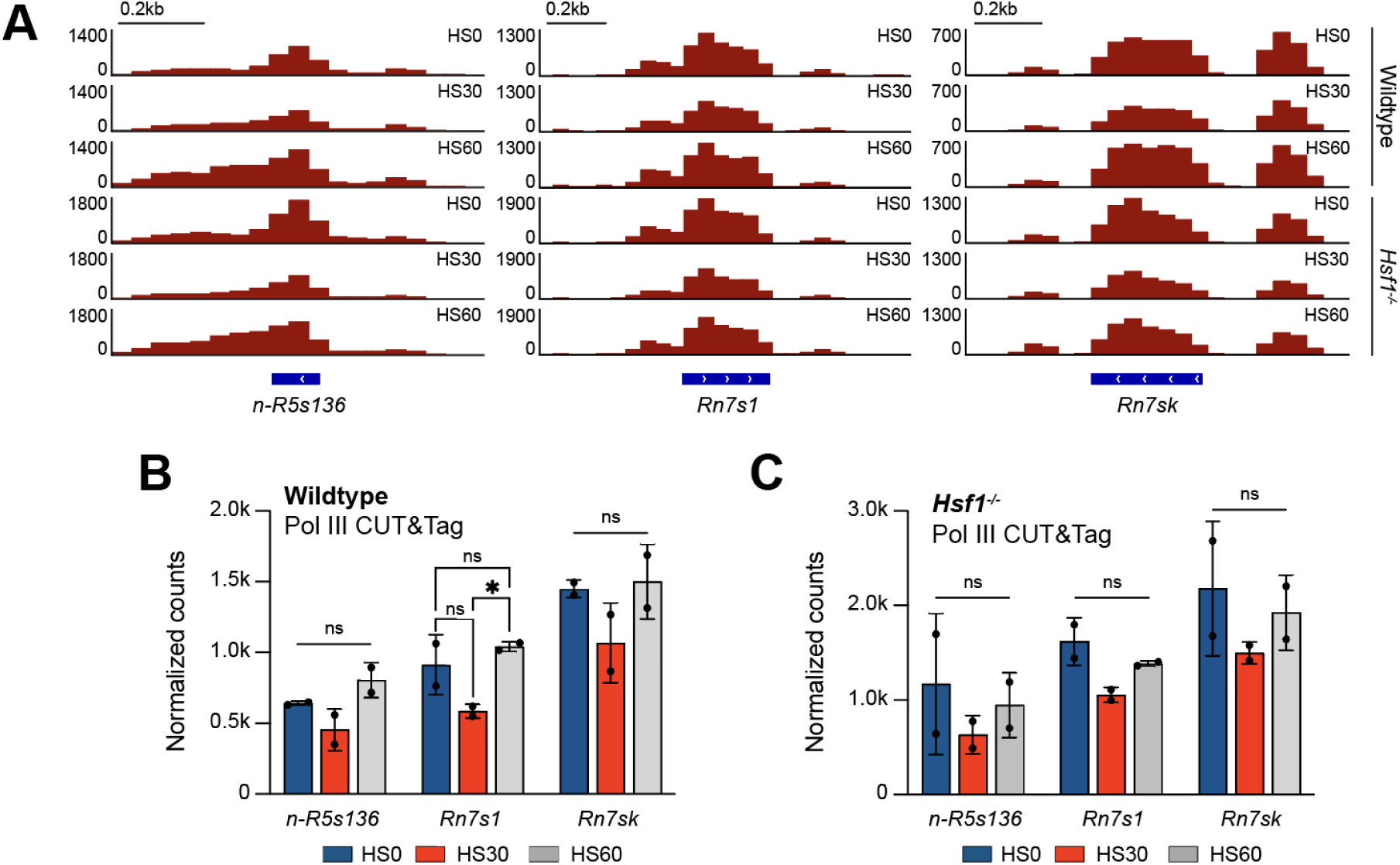
ncRNAs transcribed by Pol III are impacted during heat shock. **(A)** Gene browser tracks at *n-R5s136* (left), *Rn7s1* (middle) and *Rn7sk* (right) for reads from Pol III CUT&Tag in wildtype and *Hsf1*^-/-^ mESCs with 0 (HS0), 30, (HS30) and 60 minutes (HS60) of heat shock treatment. **(B-C)** Normalized read counts of Pol III CUT&Tag for unstressed, HS30-treated, and HS60-treated wildtype (B) and *Hsf1*^-/-^ (C) mESCs on *n-R5s136*, *Rn7s1*, and *Rn7sk* ncRNAs. CUT&Tag counts are presented as an average of two biological replicates ± SD. Statistical analysis was performed using one-way ANOVA. ns: non-significant. *: p ≤ 0.05.

We next questioned whether HSF1 played a similar role at these other classes of Pol III genes as tRNA genes. Analyzing the Pol III CUT&Tag data in unstressed, HS30-treated, and HS60-treated *Hsf1*^-/-^ mESCs at the 5S, 7S1 and 7SK genes, we observed a modest decrease in Pol III occupancy for HS30-treated *Hsf1*^-/-^ cells compared to unstressed conditions (Figure 3A, Figure S4A). However, the recovery in Pol III binding after 60 minutes of HS is blunted in the *Hsf1^-/-^* mESCs (Figure 3A, Figure S4A). These trends are further quantified in bar plot format (Figure 3C). Taken together, we conclude that these other classes of Pol III genes are regulated similarly to tRNAs during stress and that HSF1 plays a conserved role in all Pol III-transcribed genes.

### HSF1 is a determinant of heat shock-induced Pol III transcriptional memory

A well-documented component of the HSR is a mechanism that protects cells from repeated stress. The initial HS acts as a primer such that subsequent stresses elicit an accelerated response, enabling faster protection for the cell (Liu et al. 2018). This type of memory has been observed for HSF1-mediated Pol II transcription of HSP genes (Vihervaara et al. 2021), but how such protective mechanisms affect Pol III transcription is unknown. To better understand the regulation of tRNA transcription during HS memory, we performed an initial HS by exposing cells to 42°C for 60 minutes (preconditioning) followed by recovery at 37°C for 24 hours. We then performed Pol III CUT&Tag analysis after a secondary HS exposure for varying lengths of time, from no secondary stress (PHS0), to 10 (PHS10), 30 (PHS30) and 60 minutes (PHS60) (Figure 4A). In preconditioned cells, we observed that Pol III occupancy begins to be downregulated at HS10-treated cells, and that this decrease persisted in HS30– and HS60-treated cells at individual loci and at all tRNA genes (Figure 4B-C, Figure S5A). To compare the change in Pol III binding in preconditioned versus non-preconditioned samples, we calculated the log_2_ ratio of HS30 and HS60 over HS0 at all tRNA genes, and performed the same analyses for all preconditioned samples (Figure 4D). The recovery in Pol III binding after HS60 is evident in the significant difference between the log_2_ ratios of HS30/HS0 and HS60/HS0, in contrast to the lack of change in log_2_ ratios between PHS10/PHS0 and PHS60/PHS0 (Figure 4D). These findings suggest that HS memory prolongs the decrease in Pol III transcription.

**Figure 4.**
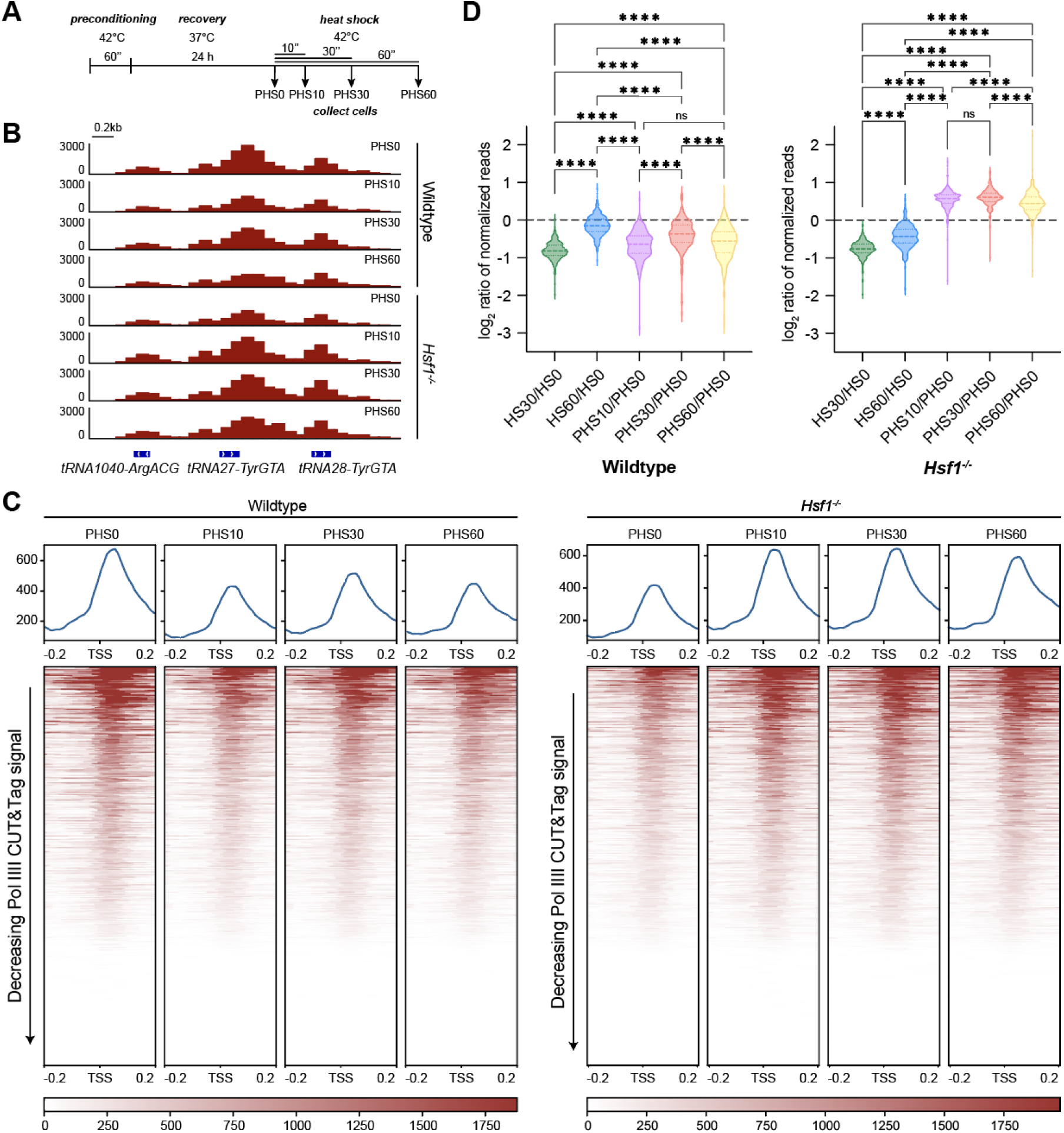
Heat shock preconditioning impacts tRNA regulation in wildtype and *Hsf1*^-/-^ mESCs. **(A)** Experimental setup for preconditioning wildtype and *Hsf1*^-/-^ mESCs. Cells were treated with a single 1-hour heat shock at 42°C, recovered for 24 hours at 37°C, and collected for Pol III CUT&Tag after an additional heat shock treatment at 42°C for 0 minutes (PHS0), 10 minutes (PHS10), 30 minutes (PHS30), or 60 minutes (PHS60). **(B)** Gene browser tracks at *tRNA1040-ArgACG, tRNA27-TyrGTA, tRNA28-TyrGTA* for reads from Pol III CUT&Tag (maroon, n=2) in preconditioned wildtype and *Hsf1*^-/-^ mESCs after 0, 10, 30, or 60 minutes of heat shock treatment. **(C)** Genome-wide average plots (top) and heatmaps (bottom) for Pol III CUT&Tag signal in a 400 bp window surrounding the TSS of all tRNA genes, arranged by decreasing signal in preconditioned wildtype (left) and *Hsf1*^-/-^ (right) mESCs after HS0, HS10, HS30, and HS60 treatments. **(D)** Violin plots depicting the fold change of normalized Pol III CUT&Tag read counts from the TSS to the TES of all tRNA genes for the listed heat shock treatments in wildtype (left) and *Hsf1*^-/-^ (right) mESCs. Statistical analysis was performed using one-way ANOVA. ns, non-significant; **** p-value ≤ 0.0001.

We then questioned the role of HSF1 on Pol III transcription of tRNAs during HS memory by performing the HS memory paradigm (Figure 4A) in *Hsf1^-/-^* mESCs. We observed that Pol III occupancy increases in PHS10 compared to PHS0 *Hsf1^-/-^* cells at individual loci and at all tRNA genes (Figure 4B-C, Figure S5B), in contrast to the pattern observed in the wild type cells. This increase in Pol III occupancy also persists for PHS30-treated and PHS60-treated *Hsf1^-/-^* mESCs (Figure 4B-C, Figure S5B). As in wild type cells, we calculated log_2_ ratio of HS over HS0 in the non– and preconditioned *Hsf1^-/-^* mESCs (Figure 4D). Without HSF1, not only does Pol III regulation become decoupled with prolonged HS, as observed by the lack of recovery in HS60/HS0, but also with transcriptional memory during repeated stress, as observed by the vastly different patterns in PHS10-PHS60/PHS0 (Figure 4D). Taken together, our results show that HSF1 plays an important role in Pol III regulation during HS and that knock-out of HSF1 leads to not only a dysfunctional HSR, but also the inability to modulate an appropriate response to recurring HS stress.

## Discussion

In this study, we demonstrate that Pol III transcription is dynamically regulated in response to heat shock (HS) stress. Specifically, we observe an initial reduction in Pol III gene expression during early HS, followed by an unexpected recovery under prolonged stress. Furthermore, we show that HSF1, the master regulator of the heat shock response (HSR), is critical for this recovery of Pol III genes during late HS and plays a role in modulating HS-induced transcriptional memory of tRNA genes.

What drives the upregulation of tRNAs during late stages of HS? Previous studies indicate that prolonged HS is accompanied by an increasing number of Pol II-upregulated genes (Mahat et al. 2016). Consequently, enhanced tRNA transcription may be necessary to meet the increased demand for protein synthesis during sustained HS. The importance of tRNAs is underscored by their depletion during mid-HS, which coincides with active translocation and degradation of tRNAs in the nucleus (Whitney et al. 2007; Miyagawa et al. 2012; Schwenzer et al. 2019). Additionally, mature tRNAs are cleaved upon HS, further depleting the tRNA pool and highlighting the need for replenishing tRNAs as the HSR progresses (Fu et al. 2009). The recovery of Pol III-transcribed tRNAs could represent an adaptive response, with Pol III activity rebounding from its HS-induced downregulation after prolonged stress (Abravaya et al. 1991). These studies suggest that the recovery of tRNA is an important step in the adaptive HSR to maintain proteostasis. Notably, this dynamic recovery contrasts with other stress responses, such as oxidative stress or nutrient deprivation, where Pol III downregulation persists over extended periods (Michels et al. 2010; Orioli et al. 2016; Torrent et al. 2018). Our findings suggest that this dynamic regulation and recovery of Pol III may be exclusive to certain stress responses, such as the HSR.

Although the role of HSF1 in Pol II-mediated HSR is well documented, its role in Pol III transcription during HS has been largely unexplored. Here, we show that HSF1 is not only important for the recovery of Pol III during late HS, but is also essential for modulating HS-induced Pol III transcriptional memory. In *Hsf1^-/-^* mESCs, we observed a significant impairment in the recovery of Pol III binding and tRNA transcription after 60 minutes of HS. This result suggests that, beyond its canonical role in regulating HSP genes, HSF1 is instrumental in the adaptive recovery of Pol III-transcribed tRNA genes during HS.

HS-induced transcriptional memory is a well-documented feature of the HSR, enabling cells to respond more rapidly to subsequent HS events. This phenomenon suggests that HS can establish transcriptional memory to accelerate protective molecular mechanisms (Liu et al. 2018; Vihervaara et al. 2021). However, the role of HSF1 in establishing HS memory for Pol III genes has been unclear. Our study shows that HSF1 plays an important role in this process. In preconditioned *Hsf1^-/-^* mESCs, tRNAs exhibited dysregulated expression during HS memory, with Pol III occupancy increasing after preconditioning rather than continuing its expected downregulation. This observation raises intriguing questions about the underlying mechanism. On one hand, HSF1 is required for Pol III recovery in non-preconditioned *Hsf1^-/-^* mESCs. On the other hand, in preconditioned *Hsf1^-/-^* mESCs, Pol III occupancy paradoxically increases after 10 minutes of HS. These contrasting findings suggest that HS attenuation and transcriptional memory are distinct yet interconnected regulatory mechanisms of the HSR, both of which appear to depend on HSF1 in complex and opposing ways. These findings underscore the intricate and dynamic nature of cellular adaptation to unfavorable environments.

## Materials and Methods

### Cell lines

For all cell lines used in this study, the parental line is JM8.N4 mouse ES cells, purchased from KOMP repository, RRID: CVCL_J962. For all experiments, a mouse *Hsf1*^-/-^ ES cell line (Price et al. 2023) or the mouse ES cell line C64 was used. The *Hsf1*^-/-^ mES cell line is a genetically modified JM8.N4 cell line containing a full deletion of the *Hsf1* gene locus as previously described (Price et al. 2023). C64 is a CRISPR-Cas9 genetically modified JM8.N4 cell line containing mAID-TBP knockin obtained as previously described (Kwan et al. 2023).

### Cell culture

Mouse ES cells were cultured on 0.1% gelatin-coated plates in mESC media: KnockOut DMEM (Corning) with 15% FBS (Cytiva HyClone), 0.1 mM MEM non-essential amino acids (Gibco), 2 mM GlutaMAX (Gibco), 0.1 mM 2-mercaptoethanol (Sigma-Aldrich), 100U/mL Penicillin (Cytiva HyClone), 100μg/mL Streptomycin (Cytiva HyClone), and 1000 units/ml of ESGRO Recombinant Mouse LIF Protein (Chemicon). Mouse ES cells were fed daily, cultured at 37°C in a 5% CO_2_ incubator, and passaged every 2 days by trypsinization. Heat shock was performed at 42°C in a 5% CO_2_ incubator for either 30 minutes or 60 minutes. For preconditioned cells, mESCs were heat shocked for 1 hour at 42°C, recovered at 37°C for 24 hours, and subjected to a single round of heat shock for 10, 30, or 60 minutes.

### Antibodies for Immunoblot

Primary antibodies: α-Tubulin 1:7000 (Abcam Cat# ab6046, RRID:AB_2210370), α-RPC7 1:2000 (Santa Cruz Biotechnology Cat# sc-21754, RRID:AB_675824), α-HSF1 1:1000 (Abcam Cat# ab2923, RRID:AB_303419). Secondary antibodies (1:15000): IRDye 800CW Goat anti-Mouse IgG (LI-COR Biosciences Cat# 926-32210, RRID:AB_621842), IRDye 680RD Goat anti-Rabbit IgG (LI-COR Biosciences, Cat# 926-68071, RRID:AB_10956166).

### CUT&Tag

CUT&Tag was performed as previously described (Kaya-Okur et al. 2019), but with the following modifications. Cells were harvested at room temperature and 100,000 mESCs were used per sample. Antibodies used include α-RPC7 (Pol III) (Santa Cruz Biotechnology Cat# sc-21754, RRID:AB_675824), Rabbit anti-Mouse IgG (Abcam Cat# ab46540, RRID:AB_2614925), and α-H3K27me3 as library positive control (Cell Signaling Technology Cat# 9733, RRID:AB_2616029). Secondary antibodies used include Guinea Pig α-Rabbit IgG (Antibodies-Online Cat# ABIN101961, RRID:AB_10775589) and Rabbit α-Mouse IgG (Abcam Cat# ab46540, RRID:AB_2614925). Secondary antibody incubation times were 1 hour at room temperature. pA-Tn5 was produced in-house by the UBC Biomedical Research Centre, and the pA-Tn5 adapter complex was added at a final concentration of 1:20. Sequencing was performed at the UBC Biomedical Research Centre using NextSeq2000 with 50 cycles.

### CUT&Tag analysis

Reads were mapped to the mm10 genome using Bowtie2 v2.4.2 (Langmead and Salzberg 2012) with the following parameters: ––no-unal ––local ––very-sensitive-local ––no-discordant ––no-mixed ––phred33 –I 10 –X 2000. The resulting SAM files were converted to BAM files using SAMtools v1.7 (Li et al. 2009). To normalize the Pol III CUT&Tag data, we used ChIP-seq-Spike-In-Free, a normalization method to determine scaling factors for samples across various conditions (Jin et al. 2020). BigWig coverage files were generated from the BAM files using deepTools v3.5.0 (Ramírez et al. 2016) and the normalization factors calculated by ChIP-seq-Spike-In-Free.

Biological replicates were merged using bigWigMerge (Kent et al. 2010), which sums the reads after normalizing each library using ChIP-seq-Spike-In-Free. Heatmaps and average plots were generated with deepTools v3.5.0, and gene plots were mapped using the Integrative Genomics Viewer (IGV) (Thorvaldsdottir et al. 2013). Read counts were generated with bedTools (Quinlan and Hall 2010), normalized with the scaling factors calculated by ChIP-seq-Spike-In-Free, and analyzed with GraphPad Prism.

### NET-seq analysis

Re-analysis of NET-seq data (GSE172401) was performed as previously described (Kwan et al. 2023). Read counts across tRNAs were generated from Drosophila spike-in normalized bam files using bedTools and analyzed with GraphPad Prism.

### PRO-seq analysis

PRO-seq data from Vihervaara et al., 2021 (GSE128160) was re-analyzed for wildtype MEFs (unstressed, HS25, HS60). PRO-seq fastq files were pre-processed and mapped to the mm10 genome using the proseq2.0 pipeline (Chu et al. 2019). BigWig coverage files were generated with BAM files using deepTools v3.5.0 and normalized by using PRO-seq reads from the 3’ end of long genes greater than 400 kb as previously described (Mahat et al. 2016). Normalized replicates were merged using bigWigMerge (Kent et al. 2010). Heatmaps and average plots were generated with deepTools v3.5.0, and gene plots were mapped using IGV.

### BRI-qPCR

BRI-qPCR was adapted from the BRIC-seq protocol published previously (Imamachi et al. 2014). The same number of cells for each sample was plated and cultured until ∼90% confluency. All cells were BrU labeled at a final concentration of 1 mM for a total of 30 minutes. For HS30-treatment, BrU was added at a final concentration of 1 mM and cells were promptly heat shocked at 42°C in a 5% CO_2_ incubator for 30 minutes. For HS60-treatment, cells were heat shocked at 42°C in a 5% CO_2_ incubator for 30 minutes, labeled with BrU (final concentration: 1 mM), and continued to be incubated at 42°C in a 5% CO_2_ incubator for 30 more minutes, totalling 60 mins of heat shock. To normalize signal across all samples, 2.5 μg of in vitro transcribed 5-BrU labeled RNA encoding for NanoLuc® Luciferase was spiked into each well, followed by Trizol extraction. BrU labeled samples were incubated at 80°C for 2 minutes, followed by immediate cooling in an iced-water bath. Protein G Dynabeads^TM^ conjugated with α-BrU antibody (MBL International Cat# MI-11-3, RRID:AB_590678) were prepared as published previously (Imamachi et al. 2014). BrU labeled samples were then incubated with the conjugated beads at 4°C for 2 hours and washed 6 times with ice-cold BSA/Triton/PBS. BrU-labeled RNAs were eluted in 90 μL 10 mM Tris–HCl at pH 7.4 and 6.25 mM EDTA and isolated via TRIzol® LS Reagent according to the manufacturer’s protocol and resuspended in 25 μL of DEPC-treated water. The entire sample was DNase treated using the Promega RQ1 RNase-Free DNase kit (M6101) and reverse transcribed using the New England BioLabs LunaScript® RT SuperMix Kit (E3010), according to the manufacturer’s protocol. The resulting cDNAs were diluted 1:5, and 4.5 μL of sample was used for each qPCR reaction with Luna® Universal qPCR Master Mix (M3003) with the QuantStudio™ 3 Real-Time PCR System.

### RT-qPCR

Cells were cultured until ∼90% confluency on tissue culture-treated plates at 37 °C in a 5% CO_2_ incubator. Heat shock was performed at 42°C in a 5% CO_2_ incubator for either 30 minutes or 60 minutes. After the indicated treatments, cells were washed with 1X PBS, trypsinized, and pelleted by centrifugation at 600 g. 2.5 μg of in vitro transcribed 5-BrU labeled RNA encoding for NanoLuc® Luciferase was spiked-in to each sample to normalize signal across all samples. RNA was extracted from the pellet by Trizol extraction and RNA concentrations were measured by Nanodrop. 1 μg of RNA was then DNAse-treated using the Promega DNAse kit (M6101), and the resulting DNAse-treated RNA was reverse transcribed using the New England BioLabs LunaScript® RT SuperMix kit (E3010). cDNA was diluted to 1 ng/μL and 4.5 ng of cDNA was used in each reaction using the New England BioLabs Luna® Universal qPCR Master Mix kit (M3003) with the QuantStudio™ 3 Real-Time PCR System.

## Data Availability

The data that support the findings of this study are openly available in Gene Expression Omnibus reference number GSE282678.

## Acknowledgments

We thank T. Stach (BRC-seq, UBC) for Illumina sequencing. This work was supported by Life Sciences Institute Cores (LSI Imaging, ubcFLOW, and qPCR Core), and by the UBC GREx Biological Resilience Initiative. For insightful comments on the manuscript, we thank Dr. Eric Jan. T.F.N., J.Z.J.K, and J.C. are supported by UBC 4-Year Fellowship, S.S.T is a Canada Research Chair.

## Funding

This work was supported by: The Canadian Institutes for Health Research Project Grant award to S.S.T. (PJT-162289); The National Sciences and Engineering Research Council Discovery Grant award to S.S.T. (RGPIN-2020-06106); The Stem Cell Network Early Career Researcher Jump Start Awards Program to S.S.T. (AWD-021244).

## Author contribution

Conceptualization: TFN, SST

Methodology: TFN, JZJK, JC, JEM

Investigation: TFN, JZJK, JC, JEM

Visualization: TFN

Funding acquisition: TFN, JZJK, JC, SST

Supervision: SST

Writing – original draft: TFN, JZJK, SST

Writing – review and editing: TFN, JZJK, SST

## Competing interests

Authors declare that they have no competing interests.

## Data and materials availability

All sequencing data have been deposited in Gene Expression Omnibus. All other data sets are available in the manuscript or in the supplementary materials. GEO IDs are in the Key Resources Table.

---

**Figure S1.**
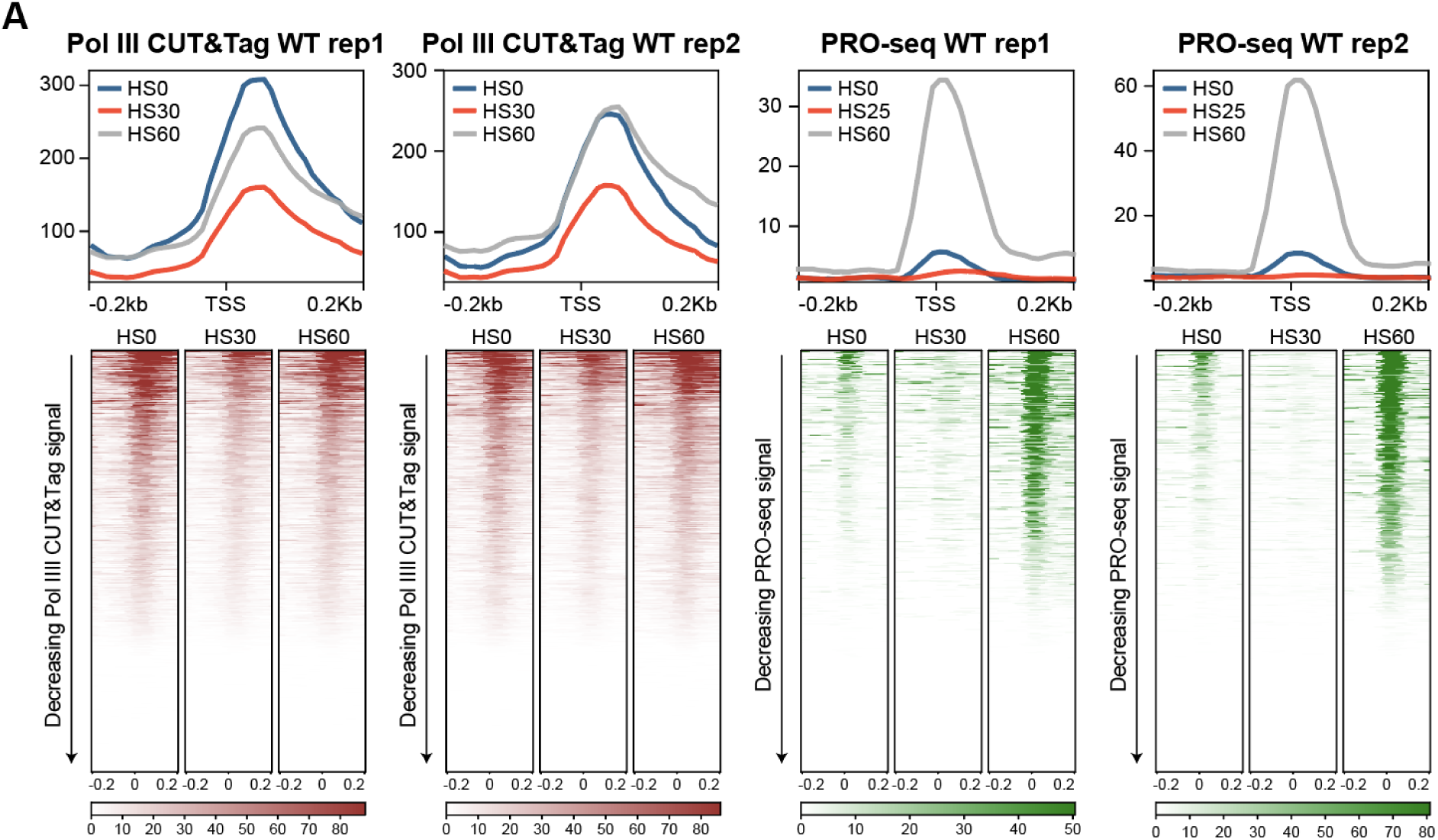
Replicate analysis of Pol III CUT&Tag in wildtype mESCs and PRO-seq in wildtype MEFs. **(A)** Genome-wide average plots (top) and heatmaps (bottom) arranged by decreasing signal for Pol III CUT&Tag (maroon) in wildtype mESCs and PRO-seq in wildtype MEFs (green) in a 400 bp window surrounding the TSS of all tRNA genes with two biological replicates.

**Figure S2.**
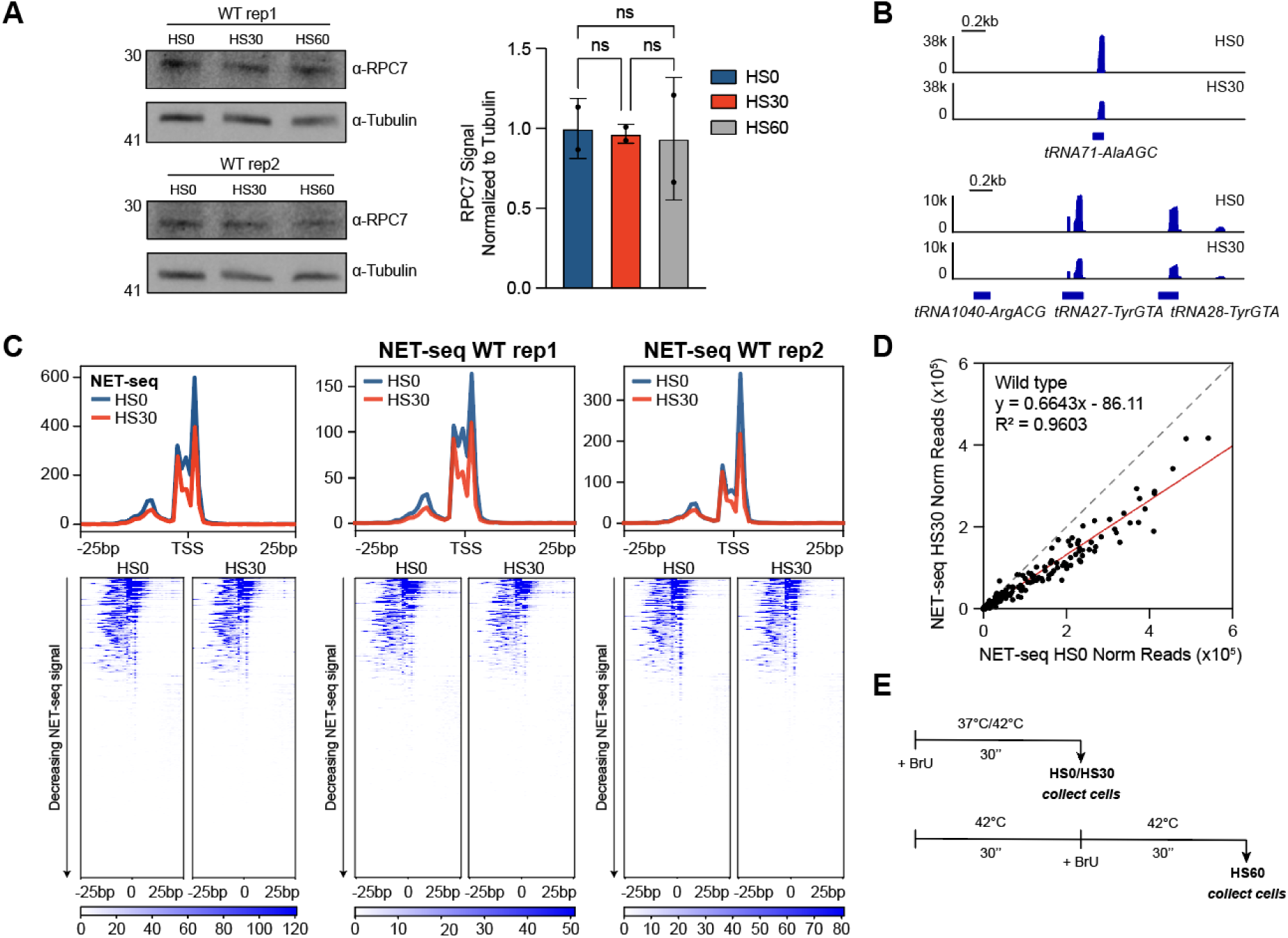
tRNA transcription is impacted upon heat shock treatment in wildtype mESCs. **(A)** Two biological replicates of immunoblot analysis (left) and relative quantification (right) with α-RPC7 (Pol III), normalized to α-Tubulin, of whole cell extracts from wildtype (WT) mESCs treated with 0 minutes (HS0), 30 minutes (HS30), and 60 minutes (HS60) of heat shock. Statistical analysis was performed using one-way ANOVA. ns: non-significant. **(B)** Gene browser tracks at *tRNA71-AlaAGC* (top) and *tRNA1040-ArgACG, tRNA27-TyrGTA, tRNA28-TyrGTA* (bottom) of reads from NET-seq in HS0 and HS30-treated wildtype mESCs. **(C)** Genome-wide average plots (top) and heatmaps (bottom) arranged by decreasing signal for NET-seq in wildtype mESCs in a 50 bp window surrounding the TSS of all tRNA genes with two biological replicates (left). Individual biological replicates are shown on the right. **(D)** Normalized read counts of NET-seq signal in HS0 vs. HS30-treated wildtype mESCs from the TSS to the TES of all tRNA genes. **(E)** Schematic for the enrichment of BrU-labeled RNA for BRI-qPCR in HS0, HS30, and HS60-treated mESCs.

**Figure S3.**
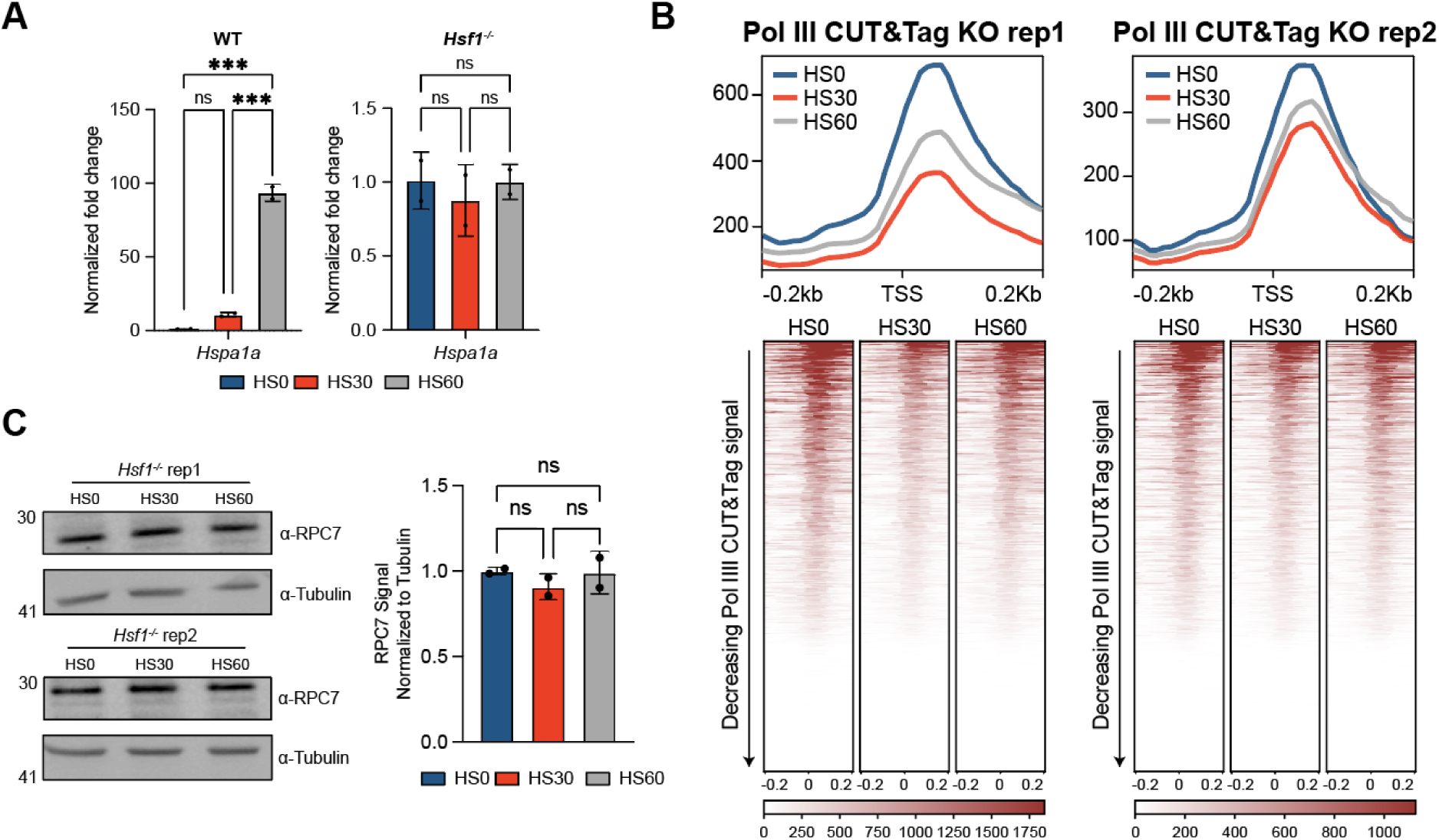
Replicate analysis of Pol III CUT&Tag on tRNAs in *Hsf1*^-/-^ mESCs. **(A)** RT-qPCR analysis of *Hspa1a* normalized to NLuc signal (n=2, mean ± SD) in wildtype (left) and *Hsf1*^-/-^ (right) mESCs treated with 0 minutes (HS0), 30 minutes (HS30), and 60 minutes (HS60) of heat shock. Statistical analysis was performed using one-way ANOVA. ns: non-significant. ***: p ≤ 0.001. **(B)** Genome-wide average plots (top) and heatmaps (bottom) arranged by decreasing signal for Pol III CUT&Tag (maroon) in *Hsf1*^-/-^ mESCs in a 400 bp window surrounding the TSS of all tRNA genes with two biological replicates. **(C)** Immunoblot analysis (top) and relative quantification (bottom) with α-RPC7 (Pol III), normalized to α-Tubulin of HS0-treated, HS30-treated, and HS60-treated *Hsf1*^-/-^ whole cell extracts. Statistical analysis was performed using one-way ANOVA. ns: non-significant.

**Figure S4.**
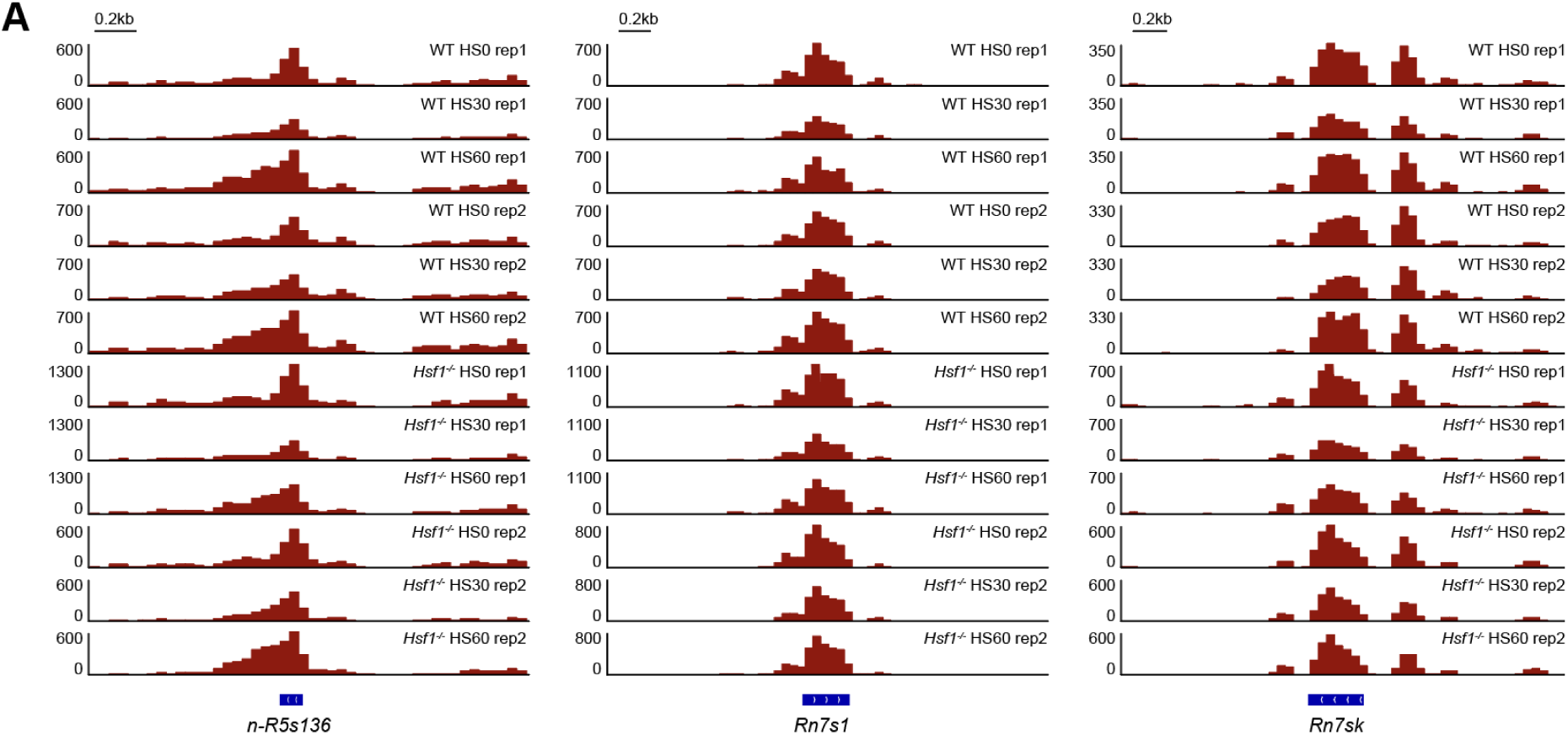
Replicate analysis of Pol III CUT&Tag on Pol III-transcribed ncRNAs in wildtype and *Hsf1*^-/-^ mESCs. (**A**) Gene browser tracks at *n-R5s136* (left), *Rn7s1* (middle), and *Rn7sk* (right) of reads from Pol III CUT&Tag in wildtype (WT) and *Hsf1*^-/-^ mESCs with 0 minutes (HS0), 30 minutes (HS30), and 60 minutes (HS60) of heat shockwith two biological replicates.

**Figure S5.**
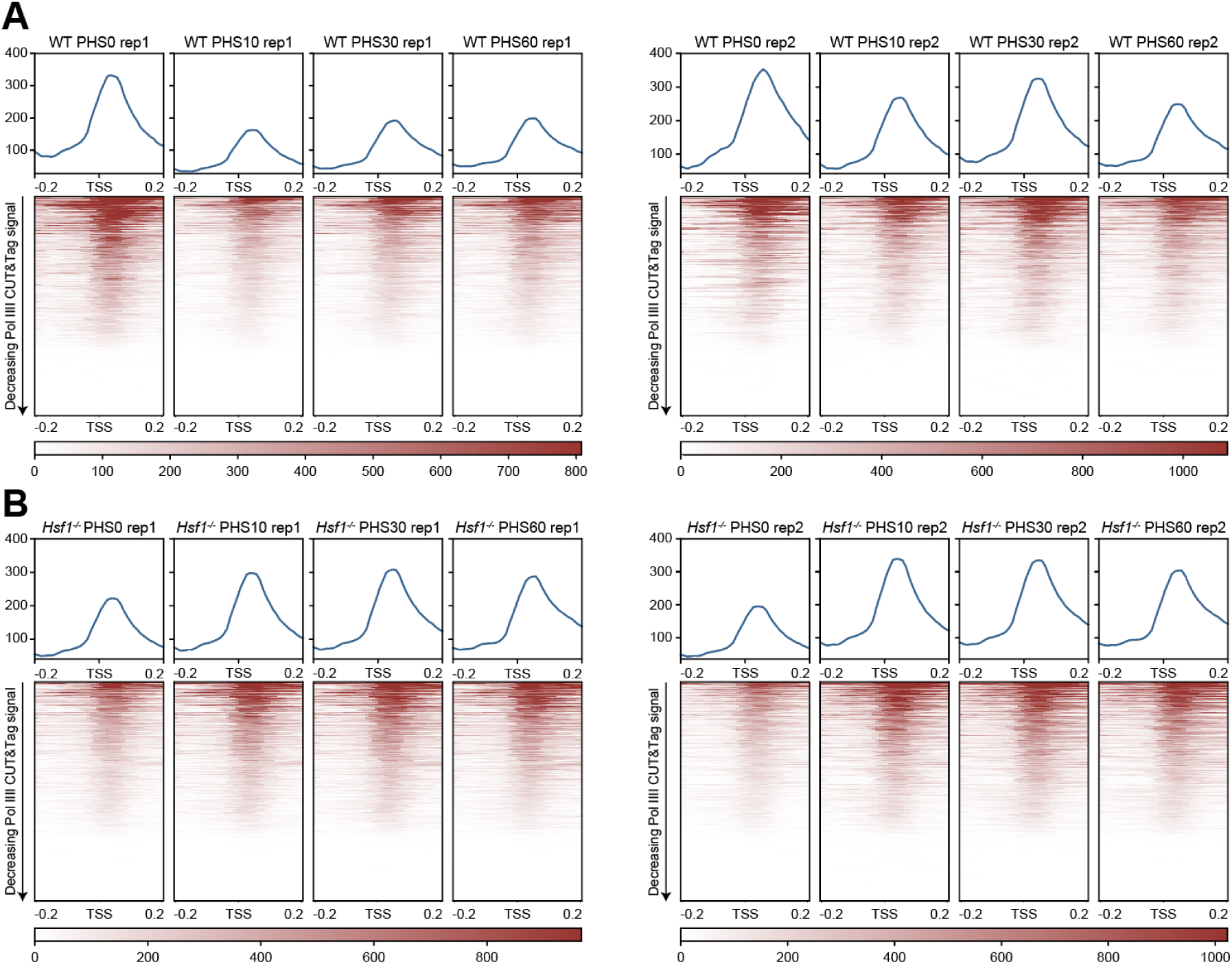
Replicate analysis of Pol III CUT&Tag on tRNAs in preconditioned wildtype and *Hsf1*^-/-^ mESCs. (**A-B**) Genome-wide average plots (top) and heatmaps (bottom) arranged by decreasing signal for Pol III CUT&Tag (maroon) in preconditioned wildtype (WT) (A) and preconditioned *Hsf1*^-/-^ (B) mESCs treated with 0, 10, 30 or 60 minutes of heat shock (PHS0, PHS10, PHS30, PHS60, respectively) in a 400 bp window surrounding the TSS of all tRNA genes with two biological replicates.

## Notes

### Competing Interest Statement

The authors have declared no competing interest.

